# Optimizing polymerase chain reaction (PCR) using machine learning

**DOI:** 10.1101/2021.08.12.455589

**Authors:** Nicholas J. Cordaro, Andrew J. Kavran, Michael Smallegan, Megan Palacio, Nickolaus Lammer, Tyler S. Brant, Vanessa DuMont, Naiara Doherty Garcia, Suzannah Miller, Tara Jourabchi, Sara L. Sawyer, Aaron Clauset

**Affiliations:** Department of Biochemistry, University of Colorado, Boulder, CO 80309, USA; Department of Computer Science, University of Colorado, Boulder, CO 80309, USA; Department of Molecular Cellular and Developmental Biology, University of Colorado, Boulder, CO, 80309, USA; BioFrontiers Institute, University of Colorado, Boulder, CO 80303, USA; Santa Fe Institute, Santa Fe, NM 87501, USA

**Author notes:** Electronic address (Aaron Clauset).

## Abstract

Despite substantial standardization, polymerase chain reaction (PCR) experiments frequently fail. Troubleshooting failed PCRs can be costly in both time and money. Using a crowdsourced data set spanning 290 real PCRs from six active research laboratories, we investigate the degree to which PCR success rates can be improved by machine learning. While human designed PCRs succeed at a rate of 55–63%, we find that a machine learning model can accurately predict reaction outcome 81% of the time. We validate this level of improvement by then using the model to guide the design and predict the outcome of 39 new PCR experiments. In addition to improving outcomes, the model identifies 15 features of PCRs that researchers did not optimize well compared to the learned model. These results suggest that PCR success rates can easily be improved by 17–26%, potentially saving millions of dollars and thousands of hours of researcher time each year across the scientific community. Other common laboratory methods may benefit from similar data-driven optimization effort.

---

Many standardized experimental protocols in biological research laboratories are time consuming and expensive to optimize. For example, polymerase chain reaction (PCR) is a particularly common technique, with commercially available DNA polymerase enzymes, optimized buffers, and vendor-provided standard protocols [1]. In a clinical setting, where a particular PCR experiment has been highly optimized and is performed over and over, e.g., the detection of SARS-CoV-2, PCR can be highly reliable. However, in a research laboratory setting PCR has many parameters that must be tuned for each new reaction [2]. These parameters include primer design, template concentration, and the temperature and time of PCR cycles, which are typically selected using a combination of scientific knowledge, past experience, and trial and error.

Reflecting the importance of PCR as a basic protocol in molecular biology research, past work has investigated various aspects of PCR design including primer creation [3], reaction setup for difficult amplification regions (amplicons) with high GC content [4], and troubleshooting for general reaction design [2, 5, 6]. Despite these efforts, novel PCR experiments often fail at nontrivial rates even in the hands of experienced researchers, and their many parameters can make troubleshooting both complex and expensive for such a basic and pervasive laboratory technique. Insight into the parameters that ultimately determine the success or failure of PCR would be indispensable in guiding researchers to better parameterize their PCR reactions, saving substantial researcher time and money.

Our data, described below, indicate that PCR experiments performed in research settings exhibit a significant failure rate of 37–45%. Considering the ubiquity of PCR across the international scientific community, such a failure rate translates into substantial waste in both time and money. To estimate the scale of this waste, we can make a rough, back of the envelope calculation as follows. According to the Carnegie Classification of institutions of higher education there are 266 research institutions in the United States with at least High research activity [7]. Assume that each institution hosts roughly 50 life science laboratories that use PCR with approximately six lab members each conducting about 30 PCR reactions per year at about $3 per reaction and 1 hour of researcher time. Hence, a 55–63% PCR success rate implies roughly $2.6-3.2 million dollars in materials cost wasted on failed PCRs and 1 million hours of time that could be reallocated to advance science.

There are now several examples of computational tools for assisting researchers in optimizing laboratory protocols. For instance, machine learning tools have recently been developed for the CRISPR-Cas genome editing system to improve activity at target sites [8] and to reduce off target affects for CRISPR-Cas assays [9–12]. Machine learning has been used to validate the selection of Threshold Cycle values for qPCR [13]. For standard PCR, work has focused on computational tools to optimize primer designs [14, 15] or to predict the PCR outcome based on primer design alone [16, 17]. These PCR tools, however, account for only a portion of the full parameter space of a PCR experiment, which limits their utility in practice.

In this study, we design a machine learning model to predict a success or failure outcome of a PCR reaction from all-encompassing input PCR parameter data, and assess its ability to improve PCR outcomes in research settings through two experiments. In the first, experimentally diverse PCR data was crowdsourced from six active research laboratories represented by 16 researchers experienced in designing PCRs for success. After transforming the recorded data into biochemically relevant features, a standard machine learning approach improved the success rate by 18–26% on held-out data. Inspecting the learned model reveals that researchers have difficulty optimizing 15 features of PCR, which our model learned to further optimize by integrating information across many different experiments. In the second experiment, we validate the models performance by using it to guide the design and predict the outcome of a new set of PCRs, by four of the same researchers across four laboratories, which simulates the way the model could be used in practice. Across these new PCRs, the models prediction accuracy was 95%, indicating a substantial real-world gain in PCR success. In our rough calculation above, improving the success rate of PCR by the 17–26% indicated by our results could potentially save $1.9 million and close to 600,000 hours of work per year.

## RESULTS

Using machine learning to optimize a laboratory protocol requires substantial and diverse data in order for the learned model to generalize. As such, data from a single biological laboratory would be insufficient, as most research laboratories utilize narrow ranges of PCR parameter values (for instance, each lab may have only one model of PCR machine). Here, we use crowdsourcing to assemble a diverse set of PCR experiments from 16 researchers across six active research laboratories, all of whom use PCR routinely. It is important to note that all of the participating researchers sought to design their PCR experiments to be successful, and hence the measured success rate represents a reasonable estimate of the baseline (general) human-expert accuracy in a research setting. Furthermore, because these experiments are highly non-random, we expect many biochemically important parameters to have little or no predictive power for success or failure, as these represent parameters that researchers are already good at optimizing. We then first apply standard machine learning techniques to integrate information across these multiple laboratories to learn a model that can accurately predict the likely success or failure of a PCR experiment, given only the experiments parameters as input. Exploring this models structure identifies which specific parameters are typically chosen in a suboptimal way. We then conduct a second validation experiment assess the models ability to improve PCR outcomes in a “live” research setting, in which four of the original researchers use the model to guide the design and predict the outcome of a set of new PCR experiments.

### Data Overview

To standardize the data collected from each researcher, we created a bespoke electronic form through which researchers recorded 37 different experiment parameters, which relate to choices about the primers, polymerase, template, and thermocycles, as well as the PCR outcome (see Materials and Methods). We restricted the data collected to standard amplification PCR, and did not collect any mutagenesis, qPCR, or colony PCR reactions. These other types of PCR can fail for reasons distinct from the reasons that a standard PCR might fail, meaning such data are unlikely to improve the machine learning models performance in our setting. Participants contributed data with a diverse set of templates ranging from human, plant, monkey, bacteria, and plasmid DNA. We note that these data represent experiments for which researchers have already tuned many reaction parameters for successful amplification and an outcome of one correctly-sized product band. As a result, they collectively represent a kind of human-expert baseline for successful PCR in a research laboratory setting. Researchers verified the outcome of each PCR by gel electrophoresis and recorded a single outcome or a combination of outcomes: (i) no bands, (ii) band(s) at the wrong size location, and (iii) correct band. In total, after dropping duplicate and incomplete records, we collected *n*_1_ = 290 PCR reactions consisting of 109 unique primer pairs with variable PCR cycle parameters and primer/template concentrations for the first experiment, and *n*_2_ = 39 additional PCR reactions for the second experiment.

### Feature Engineering

We first transformed the 37 input parameters of each of the *n*_1_ = 290 recorded reactions into a smaller set of biochemically relevant features, which maps a reaction into a 23-dimensional feature space that can be used to train the machine learning model. Using bioinformatics software packages `primer3` and `melting`, we created primer features such as homo/heterodimer and primer melting temperature (Tm). Oligonucleotide Tms are highly dependent on the free salt concentrations in solution, which is dictated by the primer, template, and dNTP concentrations [18]. To account for the free salt concentration for primer Tm prediction, we generated a library of polymerase specific buffer salt and dNTP concentrations from company specifications. All predicted Tms were then corrected using our reaction specific library of salt conditions.

A successful PCR can be defined in two ways: “clean” and “dirty.” A clean reaction can be defined as having only the PCR product at the correct band size; on the other hand, a dirty PCR contains both the correct size PCR product band and other nonspecific products at different sizes. We considered both definitions when building our model, but we focus on only clean PCR reactions for the model presented in the main text (55% of reactions). A model based on counting both clean and dirty PCRs as successes (63% of reactions) is given in the Supplemental Information (see Figs. S1 and S2). All reactions with “band at the wrong size” or “no bands” were counted as failures.

### Learning to Predict PCR Outcomes

In the first experiment, our simple goal was to learn a statistical model from our experimental data that can predict whether a PCR would be successful from its experimental parameters. To make the learned model is more interpretable, we trained a random forest classifier to predict PCR outcomes, using 1000 trees with a balanced class weight to compensate for the unequal outcomes in our dataset. The model was then validated using 10-fold cross validation averaged across six replicates with data reshuffling between replicates, resulting in 60 tests.

We then used a greedy forward feature selection algorithm to identify the most predictive subset of features without relying on Gini importance scores [19, 20]. For each possible additional feature, the model was retrained and tested using 10-fold cross validation on six replicates and the feature yielding the most positive change in predictive score was added. This process was repeated, adding features one at a time, until every feature was included in the final model. We also implemented a backward selection algorithm to evaluate whether both algorithms converged on the same most-predictive subset of features; however, the resulting model exhibited poor accuracy, likely due to overfitting, and hence no convergence was observed. One benefit of this forward selection approach is that it produces a trace of the F1 score and accuracy as a function of the sequence in which features are added, and is usually convex, with a maximum indicating the minimal and most predictive model. In our analysis, the peak in F1 score corresponds to a model with 14 features and the peak in accuracy is a model with 15 features (Fig. 1). However, we note that both are essentially equally predictive on both scores when we consider their bootstrap confidence intervals. The 15-feature model yields an accuracy of 81 ± 2%, (95% bootstrap CI) and an F1 score of 82 ± 2%, which is a 26% improvement over the baseline researcher accuracy.

**FIG. 1:**
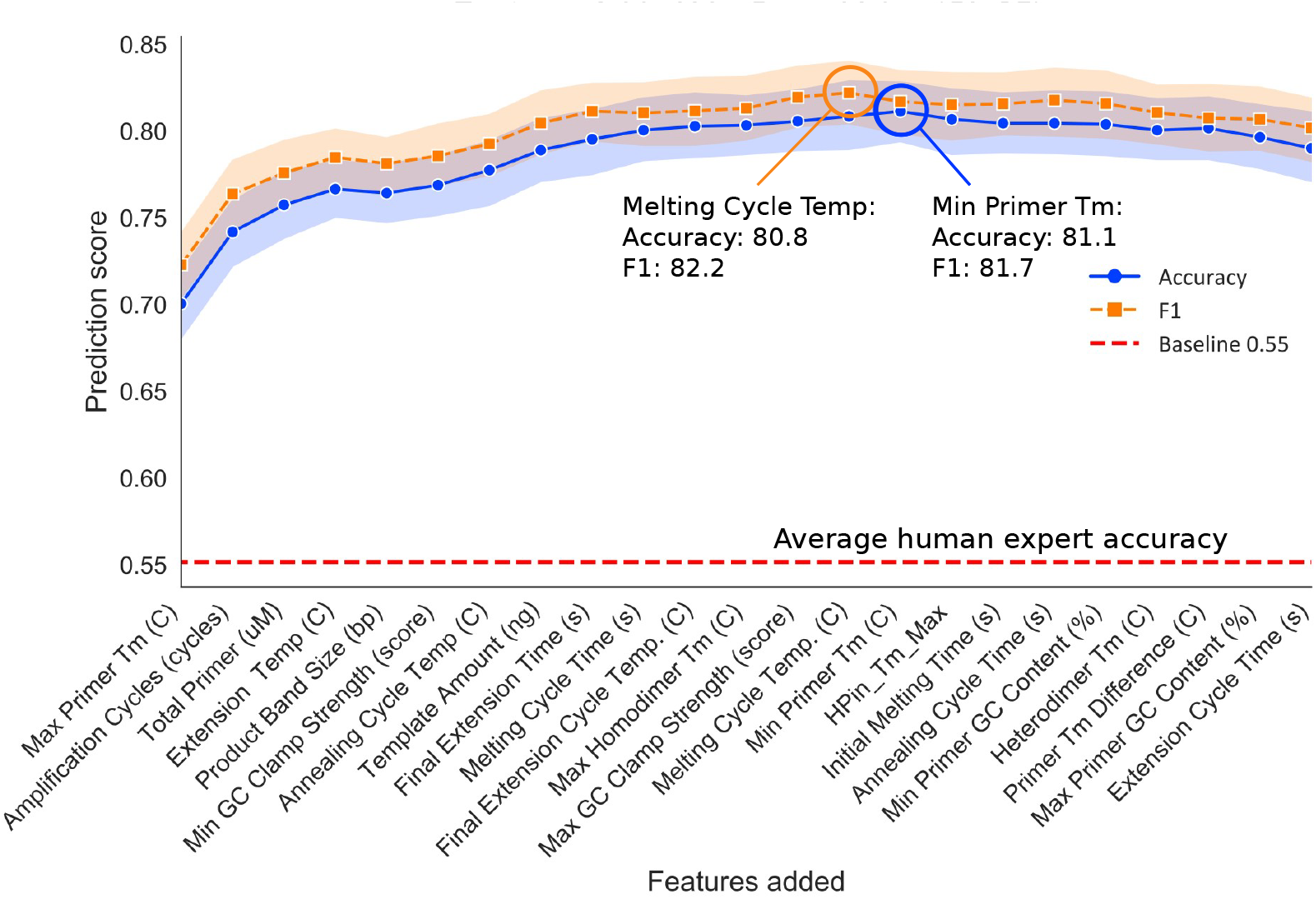
Prediction performance for the sequence of models constructed under the forward greedy selection algorithm, applied to the 23 PCR features, measured by both accuracy (fraction of predictions that were correct) and F1 score (the harmonic mean of precision and recall). Optimality is obtained at 14 or 15 features, and far exceeds the baseline accuracy of expert researchers. Envelops indicate 95% bootstrap confidence intervals, and the baseline accuracy indicates the proportion of experiments with a clean PCR outcome, i.e., the average expert researcher accuracy.

The features added after the peak, which do not improve the model, are not necessarily unimportant for PCR. Instead, the values of these features simply do not correlate with outcome in a way that can be exploited to further improve the models predictions, because researchers may have already chosen these parameters well. Similarly, the features selected prior to the maximum are not necessarily more important for successful PCR. Rather, these features represent PCR parameters that as a group researchers tend to choose suboptimally, allowing our model to improve over the researcher baseline by integrating information across many different experiments. The optimal model contains the following features: Max primer Tm, Amplification Cycles, Total Primer, Extension Temperature, Product Band Size, Minimum GC Clamp Strength, Annealing Cycle Temp, Template Amount, Final Extension Time, Melting Cycle Time, Final Extension Cycle Temp, Max Homodimer Tm, Max GC Clamp Strength, Melting Cycle Temp, and Minimum Primer Tm (Table I).

**TABLE I:** Summary statistics of 15 most-predictive features of PCR success listed in decreasing order by their contribution to model accuracy.

| features | units | %$\Delta$ accuracy | mean | SD | 25% | 75% | range |
| --- | --- | --- | --- | --- | --- | --- | --- |
| Max Primer Tm | C | 14.89 | 62.98 | 7.17 | 57.4 | 70.5 | [50.93, 81.12] |
| Amplification Cycles | cycles | 4.14 | 32.49 | 5.71 | 30.0 | 35.0 | [18.0, 44.0] |
| Total Primer | μM | 1.55 | 6.81 | 8.87 | 0.5 | 20.0 | [0.25, 20.0] |
| Extension Temp. | C | 0.92 | 71.70 | 1.23 | 72.0 | 72.0 | [65.0, 74.0] |
| Product Band Size | bp | -0.23 | 1089.95 | 1745.16 | 189.5 | 1060.0 | [2.0, 9000.0] |
| Min GC Clamp Strength | score | 0.46 | 32.68 | 4.31 | 30.0 | 35.0 | [21.0, 41.0] |
| Annealing Cycle Temp. | C | 0.86 | 59.70 | 5.75 | 55.0 | 63.0 | [50.0, 72.0] |
| Template Amount | ng | 1.15 | 335.38 | 542.73 | 1.0 | 837.7 | [0.0, 2255.6] |
| Final Extension Time | s | 0.63 | 458.07 | 130.22 | 420.0 | 600.0 | [60.0, 600.0] |
| Melting Cycle Time | s | 0.52 | 26.69 | 11.05 | 30.0 | 30.0 | [7.0, 60.0] |
| Final Extension Cycle Temp. | C | 0.23 | 71.83 | 3.43 | 72.0 | 72.0 | [65.0, 95.0] |
| Max Homodimer Tm | C | 0.06 | 11.09 | 12.59 | 0.0 | 18.8 | [0.0, 51.55] |
| Max GC Clamp Strength | score | 0.23 | 37.60 | 4.79 | 36.0 | 41.8 | [22.0, 47.0] |
| Melting Cycle Temp. | C | 0.29 | 95.49 | 1.46 | 95.0 | 95.0 | [94.0, 98.0] |
| Min Primer Tm | C | 0.29 | 58.25 | 5.66 | 54.74 | 62.5 | [46.69, 78.41] |

Table I lists summary statistics and Figure 2 depicts normalized distributions of the 15 most predictive features identified by our model. We note that the minimum observed template amount does not reflect experiments with no template. Rather it indicates researchers who were unsure about their actual template amount. Additionally, we note that a minimum product band size entry of 2bp is an entry error by a researcher. Hence, these features may be capturing differences between researchers rather than differences between reactions.

**FIG. 2:**
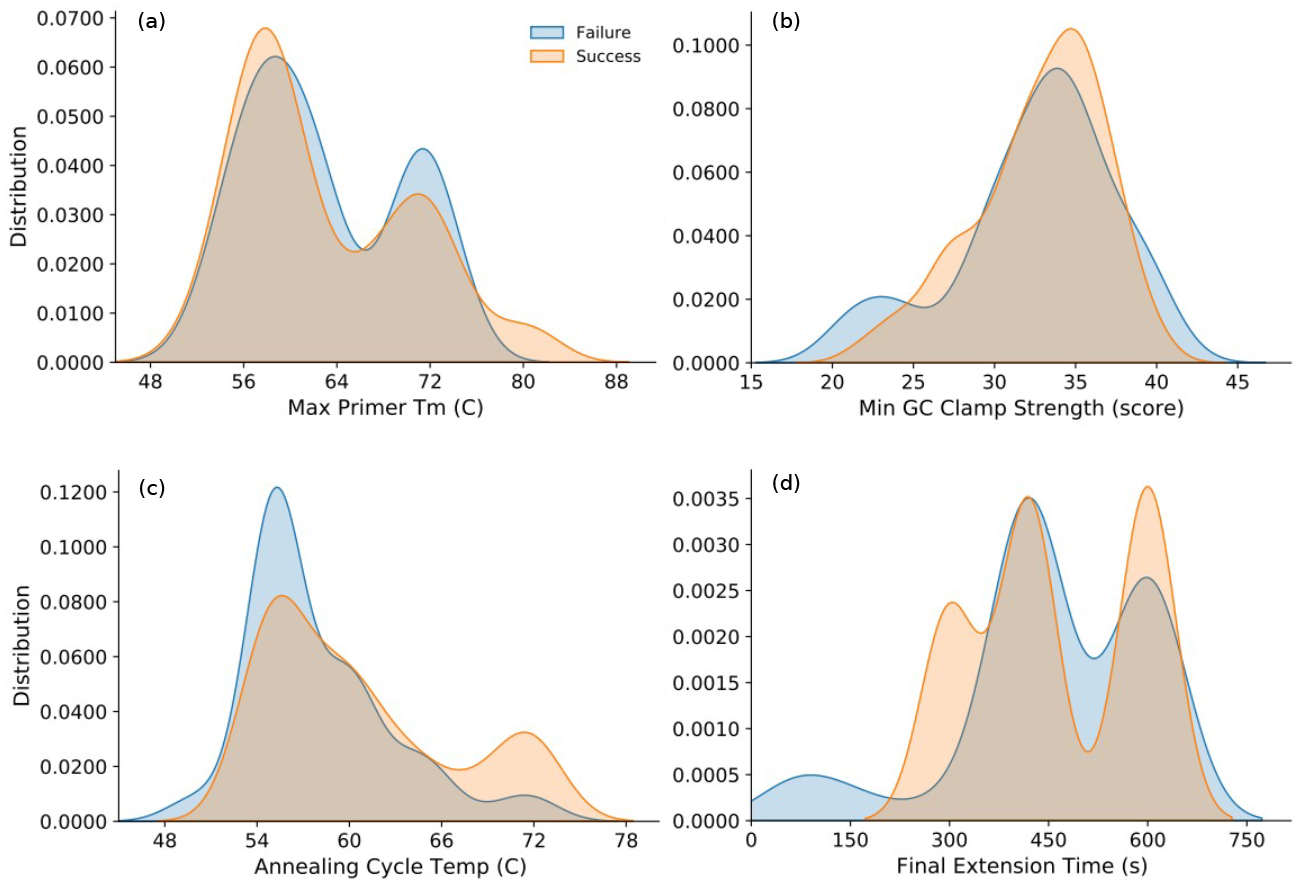
Contrasting PCR parameter distributions for success versus failure outcomes across *n*_1_ = 290 experiments, for the four most important features in the optimal prediction model (Fig. 1), (a) Max Primer Tm, (b) Min GC Clamp Strength, (c) Annealing Cycle Temp., and (d) Final Extension Time. The lack of clear separation between the contrasting distributions illustrates the marginal utility of most individual features for predicting overall PCR success, while combinations of features produce far greater prediction performance (see Fig. 1). Distributions for the remaining 11 features in the optimal model are provided in Fig. S1.

Even individual features exhibit some separation in low dimensional space, but the distinction between PCR success and failure is much larger in the higher dimensional space of the full data. Of the univariate distributions in Figure 2, those of maximum primer tm, minimum GC clamp strength, annealing cycle temperature, and final extension time are particularly notable in shedding light on how the model is optimizing success over the researcher baseline.

Maximum primer Tm (Fig. 2a), or the highest melting temperature of a primer pair for a given reaction consists of a bimodal distribution in which there is a higher proportion of successes at the lower mode and a higher proportion of failures at the higher mode. This pattern is counterintuitive because a higher primer melting temperature should help stabilize the primer template complex during amplification. The slight separation of successes and failures here may indicate an overestimation of primer Tm by researchers who, consequently, set cycle temperatures too high, destabilizing the complex.

Minimum GC clamp strength (Fig. 2b), the strength of the interaction of the 5, 3’ base pairs, has a bimodal distribution for failures, and the lower mode, located around a score of 25, has a higher proportion of failures compared to successes. This pattern suggests a threshold for GC clamp strength must be met for reaction success and GC clamp strength scores below 25 tend to fall below this threshold.

Annealing cycle temperature (Fig. 2c), the setting on the thermocycler that encourages primer template annealing following the melting cycle, contains a higher proportion of successes at higher temperatures and a lower proportion at lower temperatures. This correlation is expected because if a researcher knows their primers have a high melting temperature, they can increase their cycle parameters, making the reaction more efficient. A lower annealing temperature setting is typically set when a researcher knows their primer Tms are also low. Hence, failing to set the annealing temperature high enough may represent a common researcher error in parameterizing PCR for success.

Final extension time (Fig. 2d), the last parameter to be set on a thermocycler and which is used to clean up the reaction, contains a relatively higher density of failures at lower extension times. This correlation is also expected, because if too little time is allocated to the final clean up phase, a dirty PCR reaction will yield product containing incomplete amplicons. Our data indicates that reactions with a final extension time less than 200 seconds tend to result in failure, while reactions with a higher extension time contain fewer failures from a dirty PCR.

### Experimental Validation

In our second experiment, our goal was to assess the utility of the learned model for improving PCR outcomes in a realistic laboratory setting, in which the model is used to guide the design and predict the outcome of a new “validation set” of PCR experiments prior to the experiments being run.

Four of the same researchers from the first experiment designed and ran *n*_2_ = 39 new PCR reactions and queried the model for outcome predictions using the same parameterization as before. Reaction parameters for such queries were transformed in the same manner as in the first experiment, and the processed data was normalized using the same function as the training data in the first experiment. The resulting experiment parameterizations were used to query the trained model for success or failure predictions, and researchers were provided with these along with the biophysical properties calculated for the features in the model. Each researcher then either (i) ran the reaction with their original parameters or (ii) interacted with the model in an iterative fashion, redesigning their reaction and querying the model for its prediction until they were satisfied, prior to running the reaction. This process produced 39 new completed PCR reactions, of which 3 were redesigned once after the model initially predicted a failure outcome.

Of the *n*_2_ = 39 validation PCR reactions, 8 were predicted to fail and 31 were predicted to succeed. Of the predicted failures, 6 of 8 failed (75.0%, TN), but 2 succeeded (25.0%, FN). Of these failures, 3 had been redesigned once. Of the predicted successes, all 31 succeeded (100.0%, TP) and 0 failed (0.0%, FP). It is important to reiterate that each of these experiments was designed by a human expert with a goal of success, and the 8 experiments predicted to fail were carried out by the researchers knowing that the model had predicted they would fail. These results correspond to a prediction accuracy of 94.9%, and a combined human-and-model PCR success rate of 84.6%. This success rate accords closely with the rate we estimated via our first experiment and corresponds to an improvement of 29.4% over researcher baseline.

## DISCUSSION

Using machine learning to optimize experimental protocols in biological laboratories is a promising approach to making research more efficient by saving researcher time and funding. To investigate the feasibility of such an optimization, we focused on PCR, one of the most common lab protocols across all of molecular biology, yet it still fails in the hands of experts 37–45% of the time. Using crowdsourced data for PCR reactions, designed by researchers to succeed, we trained a simple random forest classifier to predict a PCRs outcome given only 23 input parameters. The lean model generated using our forward greedy feature selection algorithm contains 15 features. Such a model predicts the success of clean PCR reactions with an accuracy of 81 ± 2% and an F1 score of 82 ± 2%, which would lower the failure rate to only 16–20%. In a second experiment, we validated the model in a simulated “live” setting in which researchers used the model to guide the design and predict the outcome of new PCR reactions. Using the model in this way lifted the overall PCR success rate to 84.6%, in close agreement with our first experiment, indicating that PCR success rates can in practice be substantially improved by using this model to help guide the design of PCR experiments.

These results also show that, even given a relatively modest-sized data set, machine learning tools can integrate information across a diverse set of experimental settings to substantially improve protocol success in research laboratory settings. In the case of PCR, our model highlights 15 PCR experiment parameters that even expert researchers struggle to optimize: Max primer Tm, Amplification Cycles, Total Primer, Extension Temperature, Product Band Size, Minimum GC Clamp Strength, Annealing Cycle Temp, Template Amount, Final Extension Cycle Temp, Max Homodimer Tm, Max GC Clamp Strength, Melting Cycle Temp, and Minimum Primer Tm. We hasten to add that because the training data are not random, but rather experiments intended to succeed by a researcher, these 15 features do not necessarily represent the most important parameters for PCR success. Rather, they are the parameters that researchers are not already choosing well, and hence are places where even experts could improve.

The most predictive feature in the model is the maximum primer Tm and the least is minimum primer Tm. Primer Tm is important in PCR because it dictates how well a primer anneals to its template during PCRs amplification phase. We find it counter-intuitive that the max primer Tm has so much predictive power when there is no information of the second primer in the reaction at this stage in the model building. We expected the lower primer Tm to be more significant by acting as a threshold for PCR success because if one primer does not anneal well, then reaching exponential amplification is more difficult. Additionally, we expected minimum primer Tm to have more synergy with the annealing cycle temperature and extension temperature features in defining a threshold for a PCR reaction compared to maximum primer Tm. Maximum primer Tm may yield high predictive power because of the higher proportion of successes at high temperatures, or if a researcher designs one primer well they may also design the other well.

The number of amplification cycles for any given reaction was the second most predictive feature. Amplification cycles in PCR consist of a loop of primer/template melting, primer annealing, and extension, resulting in exponential amplification of the region desired in the template. If a PCR has too few amplification cycles, it may produce a product, but that product may not be abundant enough to be visualized on a gel. Additionally, if the primer annealing conditions are not ideal, more cycles may be necessary to obtain the desired amplification due to primer annealing. We also note that annealing conditions for primers are significantly influenced by total primer.

Total primer added to a PCR reaction can alter primer Tm by changing the free salt concentration in solution. Too much primer will chelate salt reducing free salt and, consequently, reduce the melting temperature because salt is required to stabilize the primer-template complex in solution. Adding too much primer can reduce the primer melting temperature [21]. Hence, if a researcher with a well-designed primer does not also account for changes in free salt concentrations, they may overestimate the true primer melting temperature in the altered environment. Consequently, they may then miscalculate their PCR parameters for the overestimated theoretical primer Tm, instead of the true primer Tm in reduced free salt. This miscalculation may explain the second mode in the maximum primer Tm distribution (Fig. 2a), where a relatively high proportion of failures occur around 72C. That is, researchers may not account for a melting temperature reduction caused by high primer concentrations and low free salt, which leads to a miscalculation of the thermocycler parameters of the melting cycle temperature, annealing cycle temperature, and extension temperature. For example, primer annealing for primers with a low Tm can be facilitated by reducing the extension temperature. However, a lower extension temperature reduces the efficiency of the DNA polymerase potentially leading to incomplete amplification of long amplicons. As a result, this feature may complement the most predictive feature, experiments primer Tm, which captures the amplicon length, band size.

The band size of the product, or the length of the amplicon, appears to contain a failed population at higher band sizes (large sections that need to be amplified). For long amplification regions, there is a greater probability of nonspecific binding, which can lead to a messy PCR reaction. Additionally, a longer amplicon will have a higher chance of containing short complementary sections that cause secondary structures potentially dislodging polymerases prior to finishing their amplification of a region. Nonspecific primer binding and secondary structures can be prevented using additives, such as DMSO. DMSO concentration was included as an input parameter for researchers during our data collection. However, almost no participants reported using it, and so this feature was dropped from the main analysis. As a result, we cannot speculate on how DMSO might alter the outcomes for large product reactions. Nevertheless, the experiment of outcome failures large products suggests that reactions with long amplicons are often parameterized sub optimally.

Minimum GC Clamp Strength and Maximum GC Clamp Strength of a primer set are both calculated using a nearest neighbor model in dinucleotide steps for the last 5 bp [22]. This 3’ clamp strength helps to “latch” the 3’ end of the primer to the template which can aid in the initiation of the polymerase. We find that a relatively high proportion of failures occurs when primers have a minimum GC clamp score less than about 25 (Fig. 2b). This pattern suggests that when the primer GC clamp is below this threshold it does not latch onto the template strongly enough for the polymerase to initiate. A low maximum GC clamp will also result in failure; however, this feature is more insignificant to the model. PCR is virtually all or nothing in regard to primers and a single faulty primer can result in reaction failure.

Reactions can also fail when primer homodimer formation Tm is high and reaction features like a lower cycle temperature permit homodimer formation, which reduces the functional primer concentration, and the concentration of primer template in the bound state. This problem is captured by a small proportion of reaction failures that occur at high homodimer temperatures (Fig. S1), which help distinguish failures when coupled with PCR cycle parameters.

Final extension cycle time influences the single cycle final reaction cleanup that allows polymerases to complete their extension and can dictate whether the PCR appears dirty or clean. Hence, this feature can help distinguish between reactions with multiple products at different bands and the correct band. However, the observed distribution of this feature (Fig. 2d) instead indicates a spurious number of failed experiments with unusually high values of this parameter. This pattern may indicate that our model has learned to exploit differences across a subset of researchers, who consistently set their final cycle time the same for each reaction.

Additional evidence that our machine learning approach is identifying same between-researcher differences can be seen in the feature template amount. Normally, this parameter can change the free salt concentration and thus potentially alter melting temperatures. However, we do not find strong evidence for this particular direct effect in our data. Instead, we find that a high proportion of successful PCRs are localized around the “0” template entry (Fig. S1), which is a special value indicating that a researcher did not know or record their template concentration.

There are several limitations to our model and data collection. Some of these limitations may serve to limit the fundamental predictability of PCR outcome by creating uncontrolled variation for the same experimental parameters, while others indicate a need for improvements in data collection for future studies of this kind. Although we adjusted all Tm estimations for the particular salt concentrations found in polymerase-dependent company buffers, the majority of these buffers have additional additives or monovalent and divalent salts differing from canonical Na^+^ and Mg^++^ used to develop Tm models. Such unlisted additives may reduce the free salt concentration resulting in an overestimation of primer Tms or may help stabilize primer hybridization leading us to underestimate the primer Tm. Additionally, we assumed the concentration of added dNTPs based on the protocols sent with specific polymerases and dNTP concentration data from researchers. Hence, unrecorded deviations from the protocols and general pipetting errors may lead to over or underestimations of Tms. As a result, we cannot correct Tm estimation errors caused by noncanonical salts or unknown additives that act in ways not captured by Tm prediction algorithms. Furthermore, the feature Template Amount, which our model found to be strongly predictive was recorded by researchers as an amount, not a concentration. Hence, varying PCR reaction volumes can significantly alter the relative concentration of template preventing standardization between reactions based on this feature. Finally, we did not collect template sequence, which precluded us from considering any features based on target effects.

We reiterate, however that the features that the models omitted are not irrelevant to successful PCRs. Instead, these parameters are likely those that researchers already optimize well, without assistance from machine learning. The fact that our model identified more than a dozen features that could be optimized further, even if a few reflect inter-researcher variability, suggests that PCR experiments are typically under-optimized, and our results provide scientifically useful hints about how to improve them.

The magnitude of the improvement in PCR success rates produced by our model is substantial, rising from the baseline of 55–63% (depending on how “clean” the success) to 81% in our cross-validation experiment and 83% in our laboratory validation experiment. With better parameterization or data collection, it may be possible to further improve this rate. For example, by collecting a larger, more representative sample of PCR experiments than those conducted by the six participating research laboratories in this study, or by using active learning techniques to efficiently collect more useful training data [23]. Even the current level of improvement, if utilized broadly, could save thousands of hours of researcher time and potentially millions of dollars in failed PCR experiments each year. Progress in this direction would only require that a researcher run their chosen PCR parameters through our model to see whether it predicts the experiment would succeed, or fail, and if the latter, then adjust the experiments parameters before moving forward, as in our validation experiment. The possibility of using machine learning to optimize other laboratory protocols, some of which are particularly complex and expensive, is an exciting direction for future work in this area.

## MATERIALS AND METHODS

We obtained data by crowdsourcing PCR reactions from 16 researchers across six active research laboratories at the University of Colorado Boulder, during the summer of 2017, and July 2019 to July 2020. Data contributors recorded PCR reaction parameters and outcomes using a bespoke spreadsheet form (Fig. 3). All data was screened for mistakes and duplicate reactions with separate outcomes were removed as user errors, e.g., forgetting to pipette a reagent. We also removed multiple replicates of the same reaction with equivalent outcomes to reduce bias in the data for any one reaction type because of our modest sample of PCR reactions. Finally, we removed reactions with primers of greater than 60 nucleotides due to the limitations of the bioinformatics algorithms that estimate melting temperature and other primer properties. This process produced *n*_1_ = 290 PCR reactions.

**FIG. 3:**
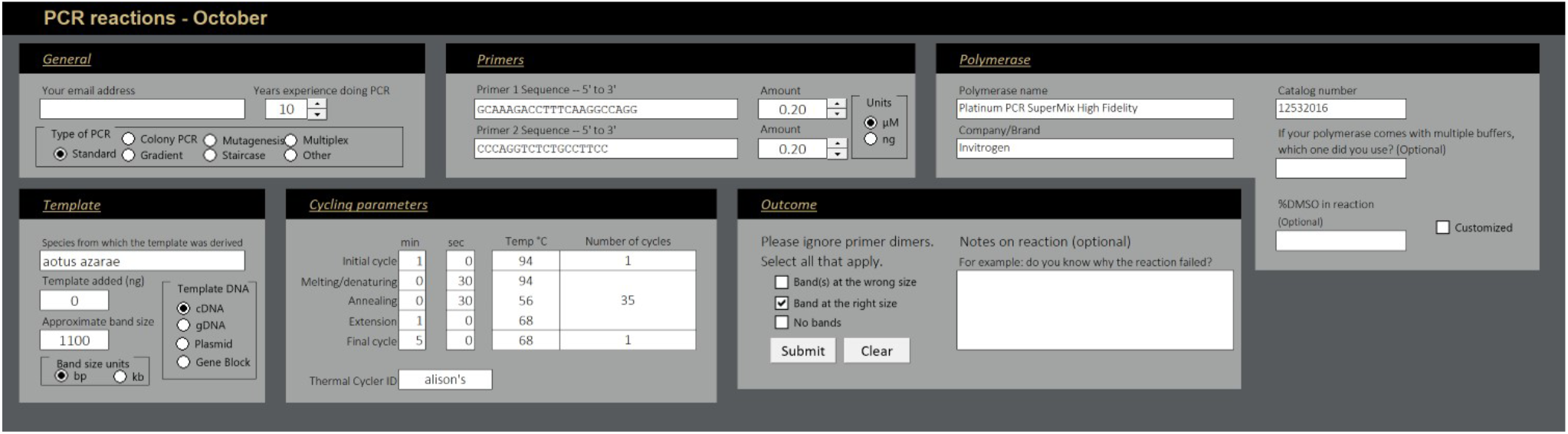
Standardized data collection form for PCR experiment parameters and outcome.

Primer properties were predicted using the python libraries `melting` and `primer3`. The library `melting` was used to predict primer melting temperatures and gave the most comparable Tm prediction to IDT, whose webtools many researchers use to predict primer Tms. The library `primer3` was used to calculate hairpin, homodimer, and heterodimer formation. Primer GC clamp strength, the 5 nucleotides on the 3’ end of the primer, was added as a feature calculated using a nearest neighbor model in dinucleotide steps for the last 5 bp [22]. The resulting GC clamp strength score considers both hydrogen bonding and base stacking interactions of the last 5 nucleotides. Features such as Tms, GC content, and clamp strength were then sorted into a maximum (Max) and minimum (Min) category per reaction to generate more interpretable boundaries for the model.

Our model was built using a random forest classifier that produced 1000 trees, was set to balanced classes, and split quality was assessed using the Gini impurity. Feature selection was carried out using a greedy forward feature selection algorithm measuring each feature addition with 10-fold cross validation across six replicates. The feature combination for each iteration with the highest accuracy was kept in the model for the next iteration.

The final model created from the first experiment, via a greedy forward feature selection algorithm and evaluated using cross validation, was then validated in a second laboratory experiment using 39 new PCR reactions, conducted between February and July 2021, and designed by 4 of the same researchers who participated in the first experiment. Each of these new PCR reactions was first designed by a researcher, and then the model-predicted outcome was returned to the researcher prior to conducting the experiment. Researchers then chose whether to (i) proceed with the experiment as designed or (ii) interact with the model iteratively before proceeding to run the experiment. Experiment parameterization and data collection was conducted in the same fashion as in the first experiment. In addition to the *n*_2_ = 39 designed and completed PCR reactions analyzed for accuracy, and additional 13 reactions were designed, but not run, by the researchers choice. Of these, 11 of 13 were predicted to fail (84.6%), and 3 of those had been redesigned once but still predicted to fail. For the 39 completed PCR reactions, a 2 *×* 2 confusion matrix tabulated the resulting combinations of predicted to fail or succeed vs. actual fail or succeed, with prediction accuracy calculated as the fraction of correction predictions out of total predictions (TN + TP)*/n*, and with success rate calculated as fraction of successful reactions out of total predictions (FP + TP)*/n*.

## Data and Code Availability

All code and data for reproducing or extending the results described in this manuscript are available at https://github.com/Oradroc/PCR_ML_model

## Acknowledgements

This research was funded in part by the Research & Innovation Seed Grant Program of the University of Colorado Boulder.

## Author Contributions

AC and SS conceptualized the research. AC acquired the funding. AK, MS, SS, and AC developed the data collection protocol. NC, AK, MS, NDG, TB, SM, TJ, VD, MP, and NL contributed data. MP, NL, TB, VD performed the validation experiments. NC, MS, and AK analyzed the data and developed the models. NC produced the figures. NC, AK, and AC wrote the initial draft of the manuscript. All authors approved the final manuscript.

## SUPPLEMENTAL INFORMATION

PCR outcome success can be defined as only having the correct amplicon, a clean outcome, or having the correct amplicon despite also having nonspecific products in the gel. Figure S2 shows the results for a second model, trained with the success parameter defined as the latter, i.e., merely containing the correct band.

The model peak accuracy contains 6 features at 79 ± 6% and an F1 score peak containing 7 features of 83 ± 4%. The features contained in this optimal model overlap with the 15 features in model with success defined as clean PCRs. We expected more features would be required to predict clean PCRs because more resolution is needed distinguish between a PCR containing the correct band alone and a correct band reaction containing nonspecific products.

---

**FIG. S1:**
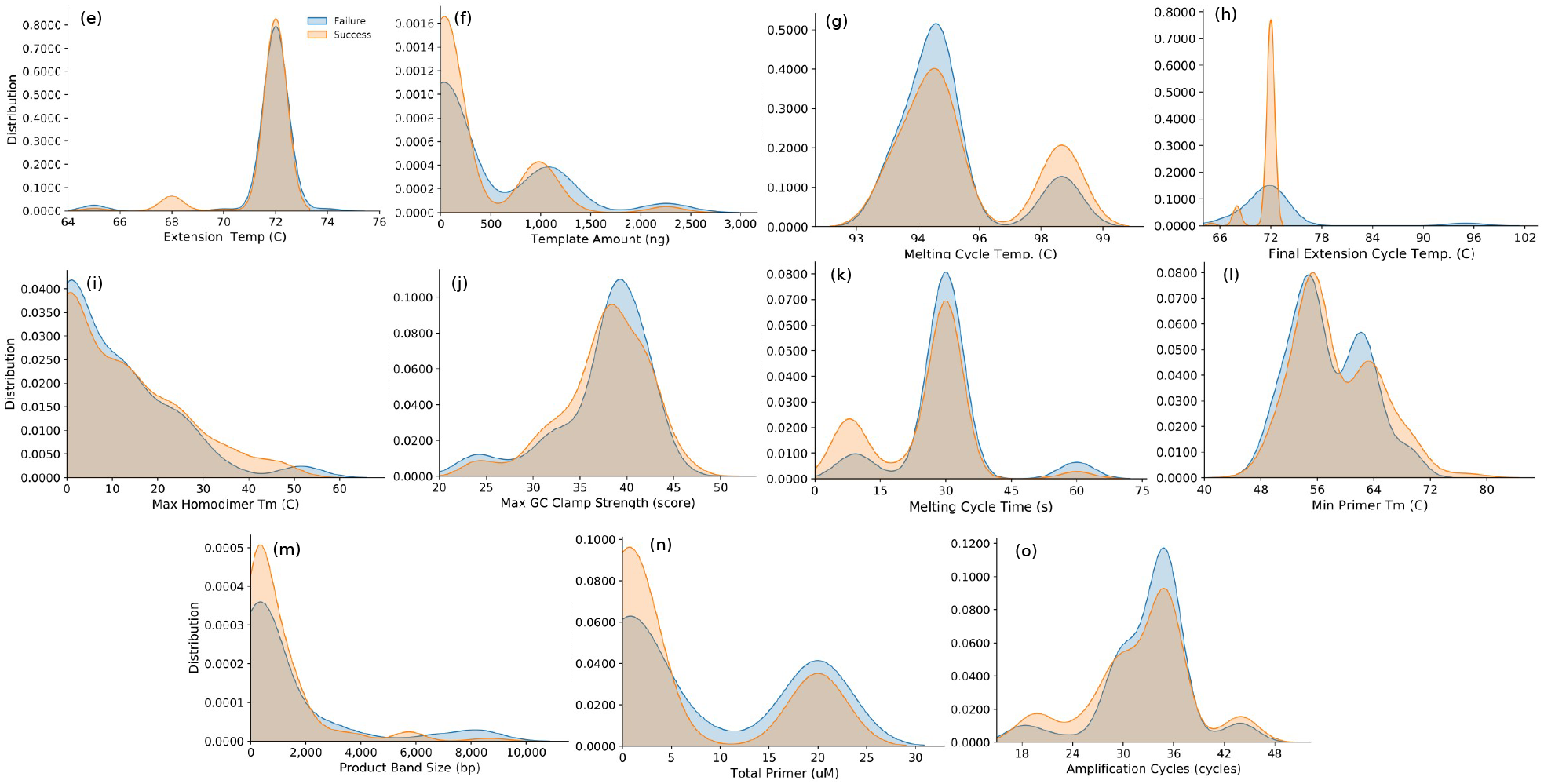
Contrasting PCR parameter distributions for success versus failure outcomes across *n*_1_ = 290 experiments, for the 11 remaining most important features in the optimal prediction model (Fig. 1), (e) Extension Temp., (f) Template Amount, (g) Melting Cycle Temp., (h) Final Extension Cycle Temp., (i) Max Homodimer Tm, (j) Max GC Clamp Strength, (k) Melting Cycle Time, (*ℓ*) Min Primer Tm, (m) Product Band Size, (n) Total Primer, and (o) Amplification Cycles. As with the four most important features (Fig. 2), the lack of clear separation between the contrasting distributions illustrates the marginal utility of most individual features for predicting overall PCR success.

**FIG. S2:**
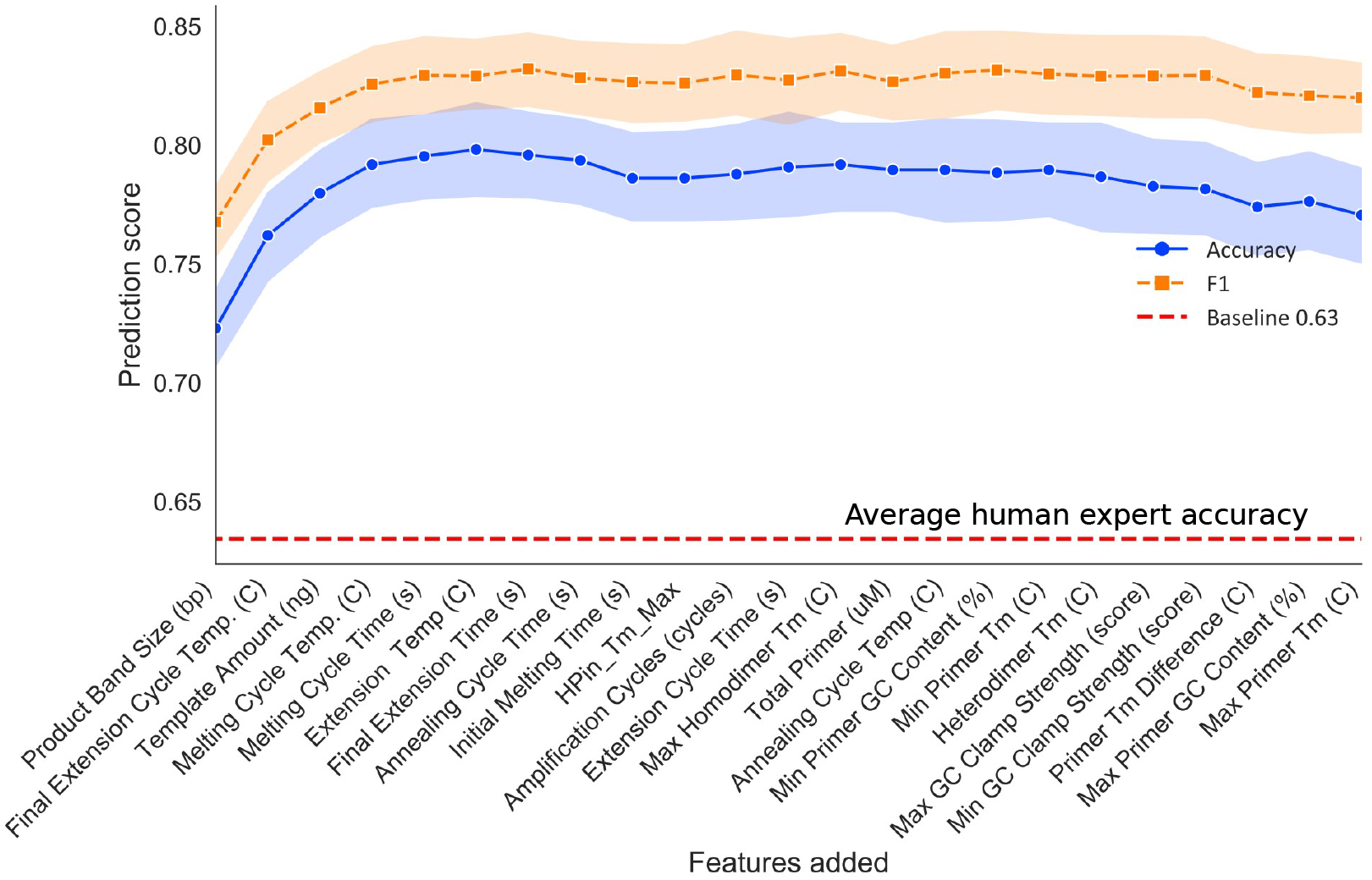
Accuracy and F1 scores for the sequence of models constructed under the forward greedy selection algorithm, applied to the 23 PCR features, with successful outcome definition allowing nonspecific products. Optimality is obtained at 6 or 7 features, and far exceeds the baseline accuracy of expert researchers. Envelops indicate 95% bootstrap confidence intervals, and the baseline accuracy indicates the proportion of experiments with a clean PCR outcome, i.e., the average expert researcher accuracy.

